# Knowledge by omission: the significance of omissions in the 5-choice serial reaction time task

**DOI:** 10.1101/2023.11.06.565747

**Authors:** Caroline Vouillac-Mendoza, Audrey Durand, Serge H. Ahmed, Karine Guillem

## Abstract

The 5-choice serial reaction time task (5-CSRTT) is commonly used to assess attention in rodents. Manipulation of this task by decreasing the light stimulus duration is often used to probe attentional capacity and causes a decrease in accuracy and an increase in omissions. However, although a decrease in response accuracy is commonly interpreted as a decrease in attention, it is more difficult to interpret an increase in omissions in terms of attentional performance. Here we present a series of experiments in rats that seeks to investigate the origins of these key behavioral measures of attention in the 5-CSRTT. After an initial training in the 5-CSRTT, rats were tested in a variable stimulus duration procedure to increase task difficulty and probe visual attentional capacity under several specific controlled conditions. We found that response accuracy reflects visuospatial sustained attentional processing, as commonly interpreted, while response omission reflects rats’ ignorance about the stimulus location, presumably due to failure to pay attention to the curved wall during its presentation. Moreover, when rats lack of relevant information, they choose not to respond instead of responding randomly. Overall, our results indicate that response accuracy and response omission thus correspond to two distinct attentional states.

## Introduction

Attention is a critical cognitive function underlying other cognitive domains, such as learning and memory, that requires careful monitoring of intermittent and unpredictable visual information over a prolonged period of time. Deficits in attentional capacities have been observed in a wide range of psychiatric disorders, including attention-deficit/hyperactivity disorder (ADHD), skizophrenia and addiction (Arnsten, 2006; Cornblatt & Malhotra, 2001; Gould, 2010; Ornstein et al., 2000). Among the number of different behavioral tasks developed in rodents to study attention and its disorders, one of the most widely used is the 5-choice serial reaction time task (5-CSRTT) (Robbins, 2002). In this task, animals must attend to a curved wall with five holes to detect a discrete stimulus light in one of them and then make a nose-poke response in the illuminated hole to receive a reward. Accurate responding thus requires that animals pay attention to a stimulus at the right time and at the right place.

Importantly, the 5-CSRTT is a flexible task that allows controlled manipulations of task parameters to test separate aspects of attention. Among these manipulations, decreasing the stimulus duration which increases task difficulty is often used to probe attentional capacity in rodent studies. Typically, this manipulation causes a decrease in response accuracy (% of correct trials) and also an increase in response omission (% of noncompleted trials) (Amitai & Markou, 2011; Asinof & Paine, 2014; Bari, Dalley, & Robbins, 2008; Robbins, 2002). An omission corresponds to a trial with no response (correct or incorrect). Decreased response accuracy is commonly interpreted as a decrease in attentional performance. However, an increase in response omission is less obvious, especially in terms of attentional performance (Robbins, 2002; Turner, Peak, & Burne, 2015). Indeed, in rodents, omitting to respond during a trial in the 5-CSRTT can occur for a number of different non-attentional factors (i.e., task-related variation in motivation).

Here we present a series of experiments in rats that seeks to investigate the origins of these key behavioral measures of attention in the 5-CSRTT. After an initial phase of habituation and training in the 5-CSRTT, rats were tested in a variable stimulus duration procedure (variable SD) to increase task difficulty and probe visual attentional capacity under several specific controlled conditions. Overall, our main results indicate that response accuracy and response omission correspond to two distinct attentional states. Response accuracy would reflect visuospatial sustained attentional processing, as commonly interpreted, while response omission would reflect rats’ ignorance about the stimulus location, presumably due to failure to pay attention to the curved wall during its presentation. Our findings also suggest that when rats lack of relevant information, they choose not to respond instead of responding randomly.

## Material and Methods

### Subjects

A total of 58 young adult male Wistar rats (300-325 g at the beginning of experiments, Charles River, Lyon, France) were used. Rats were housed in groups of 2 and were maintained in a light- (reverse light-dark cycle), humidity- (60 ± 20%) and temperature-controlled vivarium (21 ± 2°C), with water available ad libitum. Animals were food restricted (∼ 12-16 g/rat) to maintain them around 95% of their free feeding body weight. Their daily ration was given 2 hours after the end of testing sessions. All behavioral testing occurred during the dark phase of the light-dark cycle. Home cages were enriched with a nylon gnawing bone and a cardboard tunnel (Plexx BV, The Netherlands). All experiments were carried out in accordance with institutional and international standards of care and use of laboratory animals [UK Animals (Scientific Procedures) Act, 1986; and associated guidelines; the European Communities Council Directive (2010/63/UE, 22 September 2010) and the French Directives concerning the use of laboratory animals (décret 2013-118, 1 February 2013)]. The animal facility has been approved by the Committee of the Veterinary Services Gironde, agreement number B33-063-922.

### Apparatus

Height identical five-hole nose poke operant chambers (30 x 40 x 36 cm) housed in sound-insulating and ventilated cubicles were used for 5-CSRTT training and testing (Imétronic, Pessac, France) (Fig. 1a). Each chamber was equipped with stainless steel grid floors and a white house light mounted in the center of the roof. One side wall was curved inward to present an array of five circular holes (2.5 cm sides, 4 cm deep and positioned 2 cm above the grid floor), each with an internal light-emitting diode and an infrared sensor for detecting nose insertion. The opposite side wall was equipped with a delivery port with a drinking cup mounted on the midline. The delivery port was illuminated with a white light diode mounted 8.5 cm above the drinking cup. Each chamber was also equipped with a syringe pump placed outside on the top of the cubicle and connected to the drinking cup via a silastic tubing (Dow Corning Corporation, Michigan, USA).

**Figure 1.**
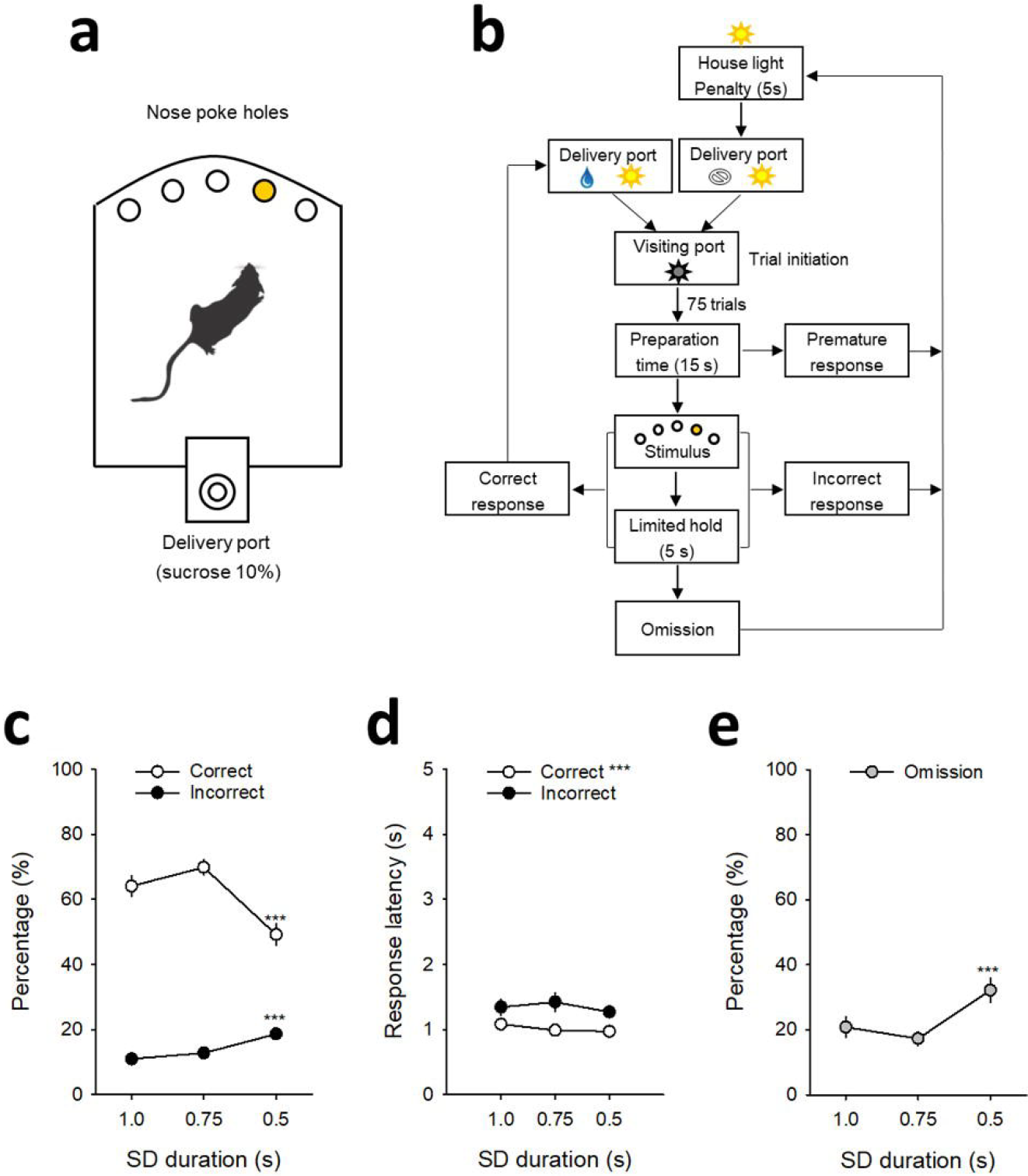
Attentional performances under the SD variable procedure. (**a**) Schematic of the 5-CSRTT apparatus and (**b**) diagram showing the sequence of events during a 5-CSRTT training session. To initiate a trial, rats had to turn off the illuminated delivery port by visiting it (i.e., exit after entering). After a fixed preparation time of 15 sec, a brief stimulus is presented in one of the 5 holes on the opposite curved wall. Animals had to nose-poke the illuminated hole (correct response), either during the stimulus presentation or within a post-stimulus limited hold (LH) period of 5 seconds, to receive a sucrose reward (0.1 ml of 10% sucrose). If they respond to the wrong hole (incorrect response), respond before the stimulus is presented (premature response), or fail to respond (omission), they are punished by a time-out penalty of 5 seconds signaled by turning on the house light. After that, the delivery port is switched on and rats have to visit it (i.e., exit after entering) to initiate the next trial. (**c, d**) Percentage (**c**) and responses latency (**d**) for correct (*white circles*) and incorrect responses (*black circles*) as a function of the light stimulus duration, respectively SD1s, 0.75s and 0.5s. (e) Percentage of omission (grey circles) as a function of the light stimulus duration (SD1s, 0.75s and 0.5s). \*\*\**p* < 0.001, different from SD1s.

**Figure 2.**
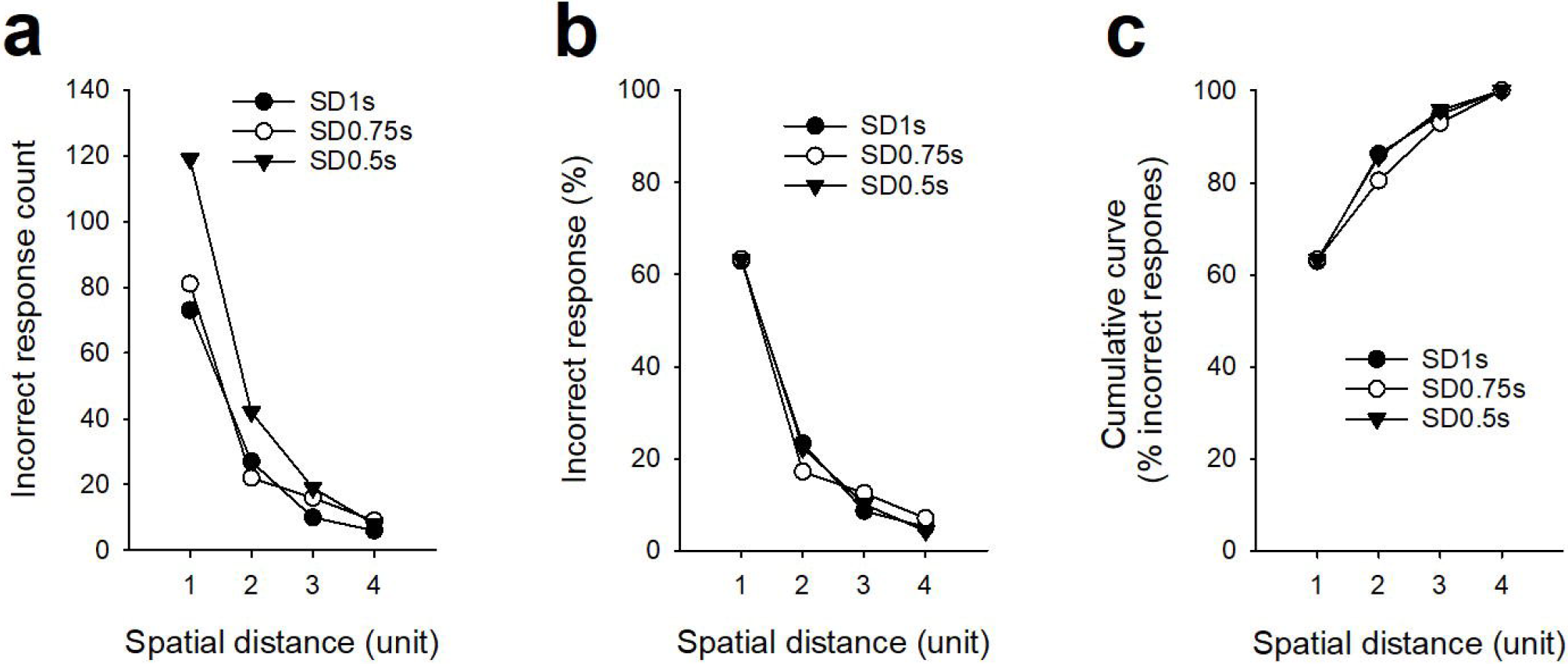
Spatial characteristic of incorrect responses. (**a**) Number, (**b**) average percentage and (**c**) cumulative curve of incorrect responses performed in a non-illuminated hole located at different spatial distance of the illuminated hole, respectively 1, 2, 3 or 4 holes next to the correct illuminated hole.

### General behavioral procedures: training and testing in the 5-CSRTT

Rats (*n* = 32) were trained with a maximum number of 75 consecutive self-paced trials per session or a maximum duration of 60 min whichever came first. The opportunity to self-initiate a trial was signaled by turning on the white light diode in the delivery port. To initiate a trial, rats had to turn off the illuminated port by visiting it (i.e., exit after entering) within one minute. If no visit occurred within the imparted time, the port was turned off automatically and this marked trial onset. After a fixed preparation time of 15 sec, a brief light stimulus was presented behind one of the 5 holes on the opposite curved wall (pseudorandom selection across trials). To receive a reward (0.1 ml of 10% sucrose), animals had to nose-poke the illuminated hole either during the stimulus presentation or within a post-stimulus limited hold (LH) period of 5 seconds (Fig. 1b). Correct nose-poke responses immediately turned off the light stimulus (if still on), turned back on the light in the delivery port and triggered the delivery of sucrose into the drinking cup. Incorrect nose-poke responses in one of the dark holes were not rewarded by sucrose, but were punished by a time-out penalty of 5 seconds signaled by turning on the house light. If animals failed to respond in any of the holes during a trial, this was considered an omission response. Omissions, like incorrect responses, were punished by a signaled 5-s time-out penalty. At the end of the time-out penalty, the house light was switched off and the light in the delivery port was turned back on for the next trial. During training, the duration of light stimulus was initially set to 30 s and progressively decreased across sessions to 1 s until the subject met performance criteria (omissions < 30%; accuracy > 60%; number of self-initiated trials > 50).

Once performance was stable with a light stimulus of 1 s (reached in 24 ± 1.2 training sessions), attentional demand was further increased within-session by decreasing the duration of the light stimulus in the following order: 1, 0.75, and 0.5 seconds. Each stimulus duration (SD) was tested during at least 25 trials per session. Animals were tested in this SD procedure during several daily sessions until stabilization of performance (reached in 15 ± 1.6 SD sessions).

### Specific experiments to probe the origin of omissions

As expected, when attentional demand increased, this caused a reduction in response accuracy, but also a large increase in response omissions. To better understand the origin of omissions, a separate group of rats (*n* = 14) was trained under the SD procedure, as described above, and then tested into specific experimental conditions. These testing conditions were designed to test the hypothesis that omissions reflect rats’ ignorance about the stimulus location, presumably due to failure to pay attention to the curved wall during its presentation (assuming that rats prefer not to respond at all than to respond randomly when they lack the relevant information). In the first testing condition, the procedure was identical to the training SD procedure, except that presentation of the light stimulus was occasionally and pseudorandomly omitted on some trials (i.e., 5 trials per stimulus duration) (Fig. 3a). There were, thus, two types of trials: normal trials with stimulus presentation (SP) and trials with stimulus omission (SO). Behavioral performances under this SO condition were assessed during 4 consecutive daily sessions. In theory, if omissions are due to ignorance, they should dramatically increase when the stimulus is effectively omitted (i.e., on SO trials in comparison to SP trials). In the second experiment, presentation of the light stimulus was signaled 1 s before by a salient change in white noise (i.e., white noise switched off and turned back on) (Fig. 4a). This was done to signal the imminence of the stimulus and, thus, to incite animals to pay greater attention to the curved wall near the time of its presentation. Behavioral performances under this signaled imminence (SI) condition were assessed during 5 consecutive daily sessions. In theory, if omissions are due to a failure to pay attention to the curved wall near the time of stimulus presentation, signaling the imminence of the stimulus should reduce omissions.

**Figure 3.**
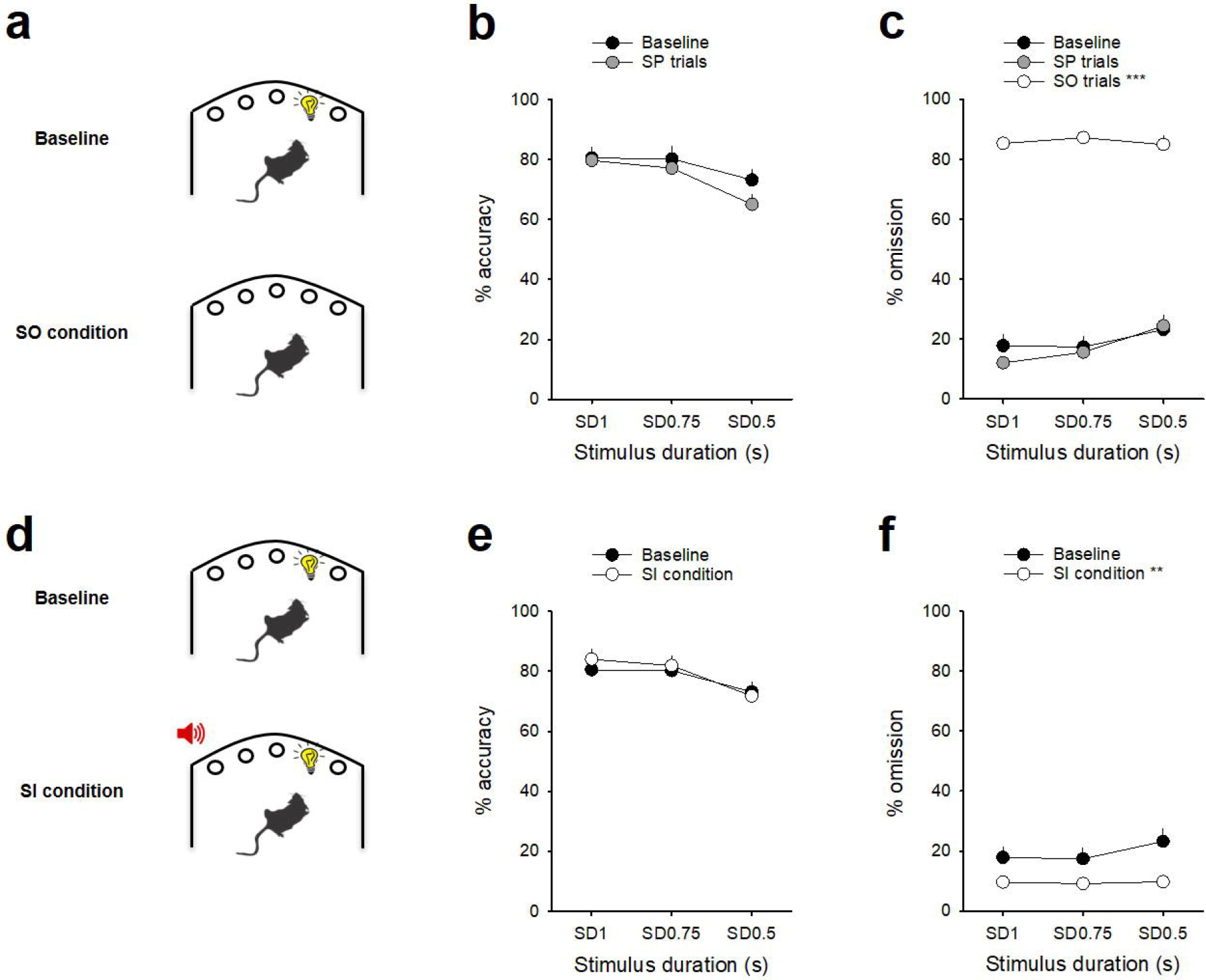
Omissions reflect rats’ ignorance about the stimulus location. (**a-c**) Animals stopped responding when light stimulus presentation was omitted. (**a**) Schematic of baseline and stimulus omission (SO) conditions in which some light stimulus presentations (i.e., 5 trials per stimulus duration) were omitted. (**b, c**) Percentage (mean ± SEM) of accuracy (**b**) and omission (**c**) during baseline (*black circles*) and during SP (stimulus presentation, grey *circles*) and SO trails (stimulus omission, *white circles*) in the SO condition as a function of the light stimulus duration (i.e., SD1s, 0.75s and 0.5s). \*\*\**p* < 0.001, different from baseline condition. (**d-f**) Animals’ performances increased when light stimulus become more predictive. (**d**) Schematic of baseline and signaled imminence (SI) conditions in which presentation of the light stimulus was signaled 1 s before by a salient change in white noise (**e, f**) Percentage (mean ± SEM) of accuracy (**e**) and omission (**f**) during baseline (*black circles*) and during the entire 5 SP conditions (*white circles*). For each session, the three circles represent the three light stimulus duration (i.e., SD1s, 0.75s and 0.5s). (**d**) Average percentage of omission (mean ± SEM) during baseline (Baseline, *black bars*) and during the last three SP conditions when light stimulus was signaled (SP condition, *white bars*) as a function of the stimulus duration (i.e., SD1s, 0.75s and 0.5s). \**p* < 0.05, \*\**p* < 0.01, and \*\*\**p* < 0.001, different from baseline condition.

**Figure 4.**
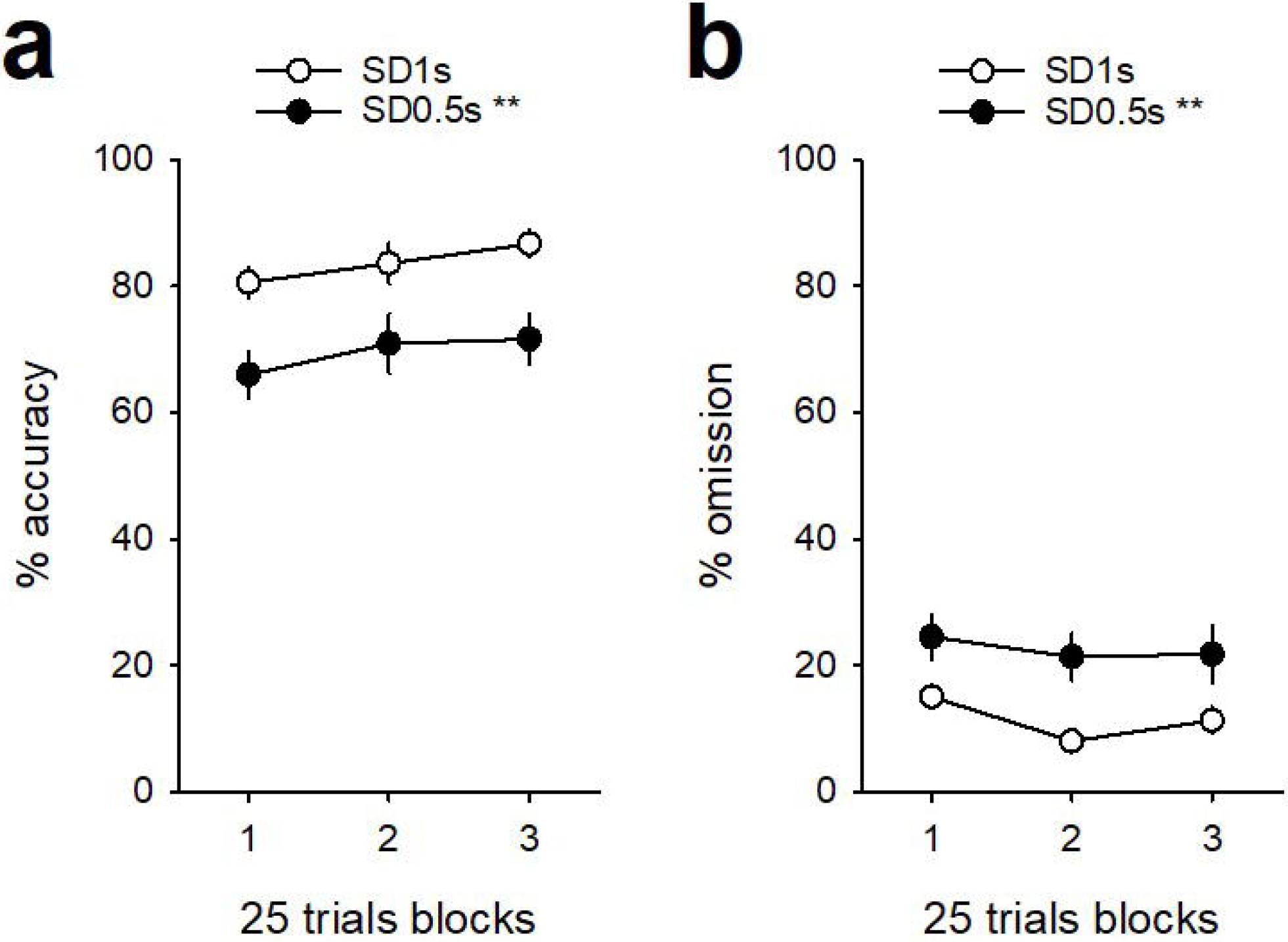
Stability of the 5-CSRTT performance within session. Within-session percentage of (**a**) accuracy and (**b**) omission during an entire SD1s (*white circles*) and SD0.5s (*black circles*) session. Each session of 75 trials was divided in 3 blocks of 25 trials. \*\**p* < 0.01, different from SD1s.

### Video Analysis

A subset of animals (*n* = 8) were also video-recorded during a SD session using an overhead camera coupled to a video tracking system (CinePlex, Plexon Inc., Dallas, TX; 30 frames/s, 1.5mm spatial resolution). This allowed us to measure animals’ head orientation and general behavior during omissions. Behavior was manually scored using a behavioral check sheet comprising: 1) head orientation toward or away from the array of 5 holes in the curved wall; 2) grooming / scratching; 3) exploration.

### Within-session measure of the degree of motivation

As the shorter SD was always tested at the end of the SD procedure (i.e., final third 25-trial block), it might be possible that the increase in omission at this particular SD might be related to a within-session decrease in the degree of motivation of the animals. To verify this possibility, another group of animals (*n* = 16) was tested with only a single SD per session (75 trials each), first with 1s, then with 0.5s. Each SD was tested for 10 days. Performance during the last 3 stable sessions for each SD were analysed across 3 blocks of 25 trials.

### Data Analysis

All data were subjected to relevant repeated measures ANOVAs, followed by Tukey post hoc tests where relevant. Statistical analyses were run using Statistica, version 7.1 (Statsoft Inc., Maisons-Alfort, France). The following behavioral measures, averaged over the last 3 stable sessions of each experimental condition, were used for analysis : % accuracy ([100 x correct responses] / [correct responses + incorrect responses]); % omission (100 x number of omissions / total self-initiated trials); premature responses (number of responses that occurred before the presentation of the light stimulus); latency of correct and incorrect responses; and, finally, reward latency (i.e., latency between correct responses and contact with the drinking cup).

## Results

Performance under the SD procedure was analysed during the last 3 stable testing sessions (Fig. 1c-e). As expected, reducing the SD caused a decrease in response accuracy, as manifested by a decrease in % correct responses (*F*_(2,62)_ = 20.7; *p* < 0.001; Fig. 1c) accompanied by an increase in % incorrect responses (*F*_(2,62)_ = 13.9; *p* < 0.001; Fig. 1c), and an increase in % omissions (*F*_(2,62)_ = 10.0; *p* < 0.001; Fig. 1e). This decrease in attentional performance was remarkably specific since it was not associated with other behavioral changes, including: latencies of correct or incorrect responses (*F*_(2,62)_ = 1.04; *NS; Table 1*), latency to collect sucrose reward (*F*_(2,62)_ = 2.6; *NS;* Table 1), number of visits in the delivery port (*F*_(2,62)_ = 1.88; *NS;* data not shown) or time spent at the drinking cup (*F*_(2,62)_ = 0.59; *NS;* data not shown).

**Table 1:**
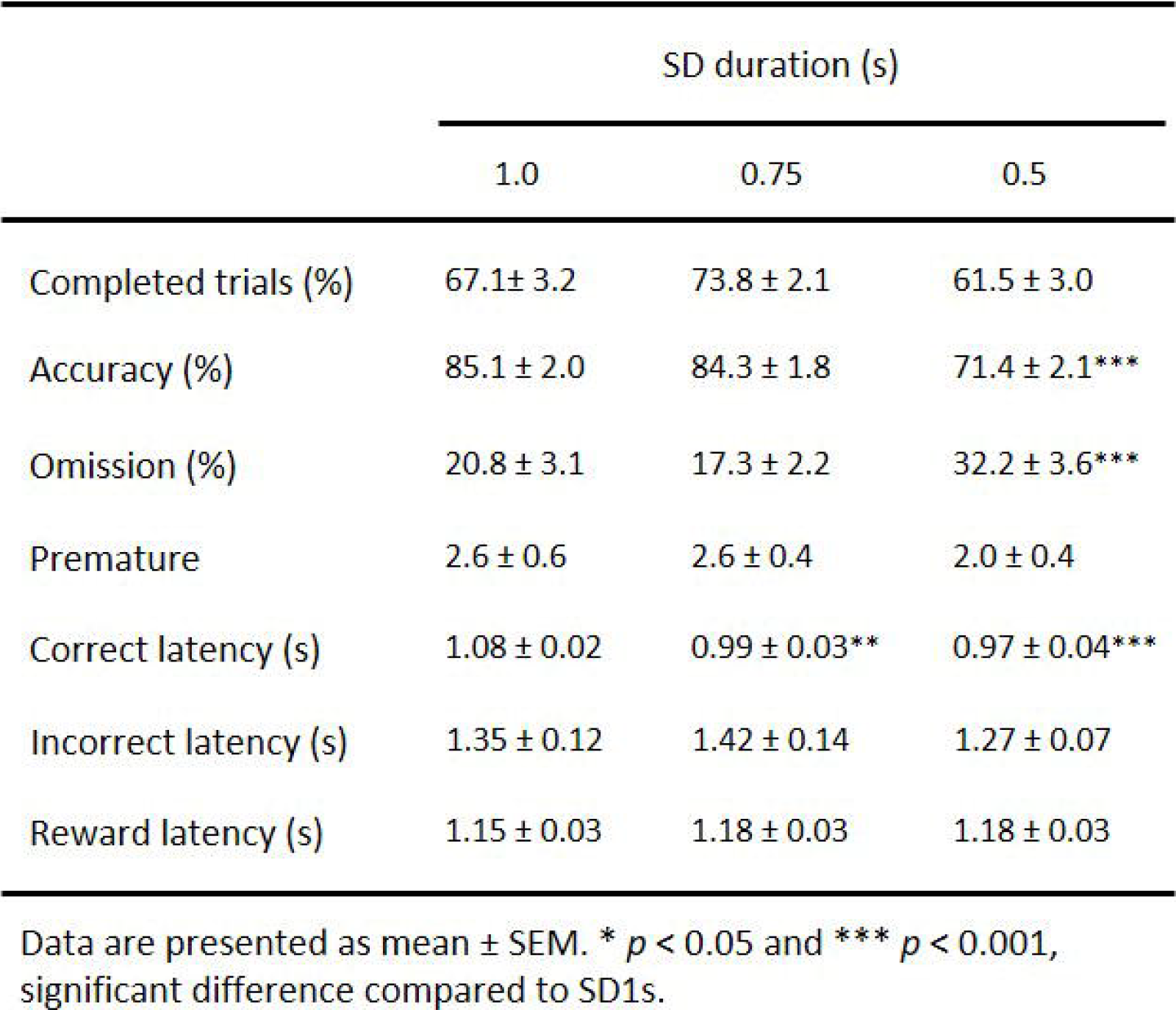
Attentional performance at SD variable.

|  | SD duration (s) |  |  |
| --- | --- | --- | --- |
|  | 1.0 | 0.75 | 0.5 |
| Completed trials (%) | 67.1 ± 3.2 | 73.8 ± 2.1 | 61.5 ± 3.0 |
| Accuracy (%) | 85.1 ± 2.0 | 84.3 ± 1.8 | 71.4 ± 2.1*** |
| Omission (%) | 20.8 ± 3.1 | 17.3 ± 2.2 | 32.2 ± 3.6*** |
| Premature | 2.6 ± 0.6 | 2.6 ± 0.4 | 2.0 ± 0.4 |
| Correct latency (s) | 1.08 ± 0.02 | 0.99 ± 0.03** | 0.97 ± 0.04*** |
| Incorrect latency (s) | 1.35 ± 0.12 | 1.42 ± 0.14 | 1.27 ± 0.07 |
| Reward latency (s) | 1.15 ± 0.03 | 1.18 ± 0.03 | 1.18 ± 0.03 |

Data are presented as mean ± SEM. \* $p < 0.05$ and \*\*\* $p < 0.001$ , significant difference compared to SD1s.

Interestingly, though the latencies of incorrect responses were slightly longer than those of correct responses (*F*_(1,31)_ = 318.2; *p* < 0.001; Fig. 1d), they were nevertheless relatively short (1.27 ± 0.07 s versus 0.97 ± 0.04 s at the shortest SD) and much shorter than the limited hold of 5 s which served as the cut-off to identify omissions. This suggests that when rats made incorrect responses, they made them with little hesitation, as if they were confident or certain, albeit erroneously, in their accuracy. This is further confirmed by the fact that, at all SD, most incorrect responses were performed in a hole very close to the correct hole (Fig. 3a-c). More precisely, 63% of incorrect responses were made in the first hole immediately adjacent to the correct hole and ∼86% of them in the first or second hole adjacent to the correct one.

Omission latencies, in contrast, were much longer (at least 5 s) than correct or incorrect responses which raises the question of their behavioral significance. In order to address this question, we trained a separate group of rats (*n* = 14) under the SD procedure, as described above, and then tested them in 2 experimental conditions: a stimulus omission (SO) condition (Fig. 3a-c) and a signaled imminence (SI) condition (Fig.3d-f) (see Materials and Methods). As expected, in the SO condition, animals made considerably more omissions on trials where the stimulus was omitted (SO trials) than on trials where the stimulus was presented (SP trials) (test condition effect: *F*_(4,52)_ = 12.41; *p* < 0.001; SO trials *vs* SP trials; p < 0.001; Fig. 3c) and this was true at all SD (test condition X SD interaction: *F*_(4,52)_ = 2.53; *NS*). Importantly, omissions or accuracy on SP trials of the SO condition were not different from baseline performance as measured before SO testing (test condition effect: *F*_(4,52)_ = 12.41; *p* < 0.001; SP trials *vs* baseline; *NS*; Fig. 3c and *F*_(2,26)_ = 1.51; *NS*; Fig. 3b, for omission and accuracy, respectively). Conversely, in the SI condition where the imminence of the stimulus presentation was signaled by a change in ambient white noise 1 s before, animals made fewer omissions in comparison to baseline performance as measured before SI testing (*F*_(1,13)_ = 13.3; *p* < 0.01; Fig. 3f). In contrast, this manipulation had no effect on accuracy (*F*_(1,13)_ = 0.54; *NS*; Fig. 3e).

Together these results indicate that omissions likely reflect rats’ failure to pay attention to the curved wall at or near the time of the presentation of the stimulus light. To verify this, we measured and quantified animals head orientation and general behavior during omission trials at the shorter SD duration (i.e., 0.5 s), using video recordings in a subset of rats (*n* = 8). As expected, during omissions, the vast majority of the animals (67 % ± 7) were not looking straight at the 5 holes and had their head turned away from the curved wall during the onset of the light stimulus presentation (data not shown). Moreover, rats were also highly engaged in another behavior in 72 % ± 8 of the time (grooming / scratching 81 % ± 10 and exploration 19 % ± 10) during the presentation of the light stimulus during omissions. Importantly, the increase in omissions seen at the shorter SD was not due to a within-session decrease in motivation as no significant change in performance was observed across blocks of 25 trials when rats were tested with the same single SD (1 or 0.5s) during the whole session (Fig. 4; see Materials and Methods) (SD X block interaction: *F*_(2,30)_ = 0.17; *NS*; Fig. 4a and *F*_(2,30)_ = 0.54; *NS*; Fig. 4b, for accuracy and omission, respectively).

Since animals are less accurate and make more omissions when the SD lasts 0.5s versus 1s, this may suggest that they pay less attention to the curved wall at the shorter SD. This was not the case, however. More animals had their head turned away from the curved wall during omission trials when the SD lasted 1s than when it lasted 0.5s (89 % ± 7 and 67 % ± 7, respectively; *F*_(1,7)_ = 7.12; *p* < 0.05; data not shown). Similarly and perhaps more surprisingly, more animals looked away from the curved wall when the SD lasted 1s than when it lasted 0.5s on correct trials (26 % ± 3 and 13 % ± 3, respectively; *F*_(1,7)_ = 7.12; *p* < 0.01; data not shown). Overall, these findings may suggest that animals allocate more attention to the curved wall at the shorter SD presumably in an attempt to maximize their chance to detect the stimulus light.

Finally, we tested whether omissions were related to individual differences in animals’ performance in the 5-CSRTT. When the SD decreased from 1s to 0.5s, the vast majority of rats decreased their accuracy (individual range: 50%-100% at 1s to 52.1%-19.2% at 0.5s; Fig. 5a), and most individual delta in accuracy were below 0 (range: -52.1% to 19.2%; mean: -13.7 ± 2.3%; median: -14%; Fig. 5b). Animals were then ranked according to their individual delta in accuracy and split by the median into low (LP) and high performers (HP). When the SD decreased, LP rats did not maintain their performance and exhibited a decrease in delta accuracy larger than that in HP rats (*F*_(1,30)_ = 31.80; *p* < 0.001; Fig. 5c). LP rats also displayed a larger increase in delta omission than HP rats (*F*_(1,30)_ = 6.52; *p* < 0.01; Fig. 5d). Importantly, these changes in performance when the SD decreased were not due to initial differences between the two groups (SD1s: *F*_(1,30)_ = 1.97, *NS* and *F*_(1,30)_ = 1.60, *NS* for accuracy and omission, respectively; data not shown). Finally, changes in delta accuracy were negatively correlated with changes in delta omission (*r* = -048; *p* < 0.01; Fig. 5e) indicating that these two variables covary when the attentional task becomes more difficult.

**Figure 5.**
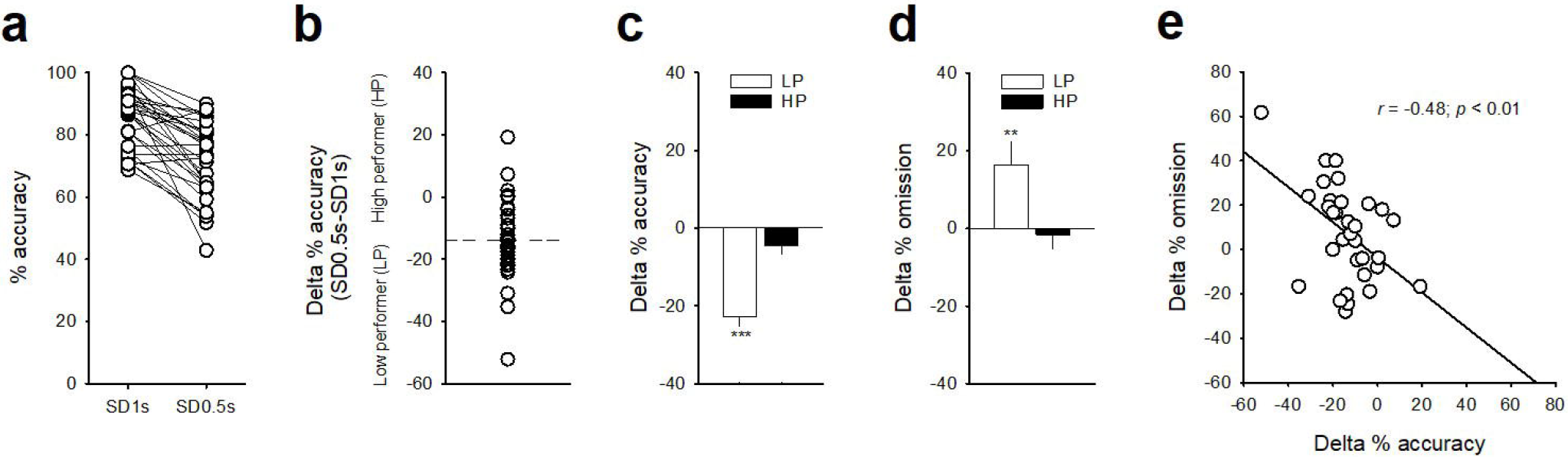
Individual differences in 5-CSRTT performance. Distribution of individual percent score of (**a**) accuracy during SD1s and SD0.5s and (**b**) delta in accuracy (i.e., SD1s - SD0.5s). Animals were then split by the median into low (LP) and high (HP) performers rats. (**c, d**) Mean (± SEM) delta in accuracy (**b**) and delta in omission (**c**) in low (LP; white bars) and high performers rats (HP; black bars). \*\**p <* 0.01 and \*\*\**p* < 0.001, different from HP rats. Correlation between the delta in accuracy and the delta in omission.

## Discussion

This study aimed to disentangle the nature of key behavioral measures reflecting attention in the 5-CSRTT. Consistent with previous findings (Amitai & Markou, 2011; Bari et al., 2008; Dalley et al., 2005; Mirza & Stolerman, 1998; Robbins, 2002), decreasing the light stimulus duration from 1 to 0.5 s was associated with a decrease in response accuracy (% of correct trials) and a large increase in response omission (% of noncompleted trials). Importantly, these effects were relatively selective since virtually all other behavioral measures provided by the task, such as premature responses, response latencies and reward latencies, remained unchanged. Thus, decreasing the light stimulus duration appeared to selectively tax attentional function, without affecting other aspects of 5-CSRTT performance, such as, for instance, response inhibition or speed.

The fact that response accuracy decreased with no or little change response latencies is intriguing and suggests that rats were equally confident in their decision between correct and incorrect responses. There is indeed evidence that response latencies are related to the degree of decision confidence (Fetsch, Kiani, & Shadlen, 2014; Kiani, Corthell, & Shadlen, 2014). Thus, our results may suggest that when rats made incorrect responses, they made them as if they were confident that they were making correct responses. Consistent with this idea, further analyses showed that at all SD, most incorrect responses were performed in a hole very close to the correct hole. More precisely, 63% of incorrect responses were made in the first hole immediately adjacent to the correct hole and ∼86% of them in the first or second hole adjacent to the correct one.

In contrast, omission latencies were discontinuously much larger than either correct or incorrect responses, particularly at the shorter SD where omissions were the most frequent. This may suggest that response omissions reflect rats’ ignorance of the stimulus location. Indeed, being ignorant of where the stimulus was presented, rats should remain undecided for some time until the next trial opportunity or before eventually responding randomly if such opportunity was too long (which was unlikely to be the case in our study because of its short limited hold of 5s). To address this question, we conducted a series of specific observations and experiments. First, we confirmed by direct observation that on omission trials, animals were not paying attention since their head and body were turned away from the curved wall at the onset or near the time of the presentation of the stimulus light, thereby preventing them from obtaining the required information about its location. Second, we confirmed that when the light stimulus was effectively omitted on some trials, thereby causing a known state of ignorance, this dramatically increased response omissions. Finally, when the imminence of the light stimulus was signaled, thereby inciting animals to pay attention to the curved wall, this reduced response omissions. Overall, our work strongly suggests that response omissions reflect rats’ ignorance about the stimulus location, presumably due to failure to pay attention to the curved wall during its presentation. This conclusion is consistent with previous observation showing that well-trained rats will often withhold a response rather than ‘guess’ when they are uncertain about which hole was illuminated (Amitai & Markou, 2011; Asinof & Paine, 2014; Robbins, 2002).

It is important here to stress that this failure caused increased omissions only when the SD lasted 0.5s, but not when it lasted 1 s. This suggests that when the SD is sufficiently long, rats may on many occasions have enough time to reorient their head toward the curved wall to accumulate sufficient visual information about its location to respond. This would not be the case, at least on most occasions, when the light stimulus is too short (i.e., 0.5s), thereby leaving rats in a state of ignorance about its location. This interpretation is consistent with what we know about how the brain makes perceptual decisions based on time-dependent accumulation of sensory evidence (Gold & Shadlen, 2007; Lo & Wang, 2006; Smith & Ratcliff, 2004). Moreover, as the accumulation rate is proportional to the amount of the visual evidence, response latencies, as indices for the time to decisional threshold, increase with decreasing sensory evidence and should thus be longer when the SD lasts 0.5s than when it last 1s which corresponds to what was observed in the present study.

While attention is most commonly measured by accuracy of responding (Amitai & Markou, 2011; Bari et al., 2008; Robbins, 2002), our results indicate that omissions also reflect a lack of enough visual accumulation due to rats’ failure to pay attention to the curved wall during light stimulus presentation. This is particularly relevant as omission rates are an important measure on human continuous performance task (Conners, Epstein, Angold, & Klaric, 2003) and have recently been shown to represent endophenotypes in ADHD (Acosta-Lopez et al., 2021). Moreover, because response accuracy and omission reflect different aspects of attention it will be interesting in future research to see whether these two behavioral parameters rely on distinct brain regions and/or neurophysiological mechanisms.

## Author contributions

K.G. and S.H.A designed research and experiments; C.V.M, A.D. and K.G. performed behavioral experiments and associated data analysis; K.G. and S.H.A. wrote the paper.

